# Diverse tuning underlies sparse activity in L2/3 vibrissal cortex of awake mice

**DOI:** 10.1101/457796

**Authors:** Yadollah Ranjbar-Slamloo, Ehsan Arabzadeh

**Author notes:** Corresponding author: Ehsan Arabzadeh.

## Abstract

It is widely reported that superficial layers of the somatosensory cortex exhibit sparse firing. This sparseness could reflect weak feedforward sensory inputs that are not sufficient to generate action potentials in these layers. Alternatively, sparseness might reflect tuning to unknown or higher-level complex features that are not fully explored in the stimulus space. Here, we examined these hypotheses by applying a range of vibrotactile and manual vibrissal stimuli in awake, head-fixed mice while performing loose-seal cell-attached recordings from the vibrissal primary somatosensory (vS1) cortex. A high-velocity stimulus delivered by a piezo-electric actuator evoked activity in a small fraction of regular spiking supragranular neurons (29%) in awake condition. However, a majority of the supragranular regular spiking neurons (84%) were driven by manual stimulation of whiskers. Our results suggest that most neurons in the superficial layers of vS1 cortex contribute to coding in awake conditions when neurons may encounter their preferred feature(s) during whisker-object interactions.

## Introduction

Spiking activity is the unit of communication across cortical neurons and is key to encoding the external world and in generating sensory perception (Barlow 1972). In a classic single unit recording experiment, Dykes and Lamour found a surprisingly high number of silent neurons in the somatosensory cortex; only a quarter of neurons recorded across cortical layers were responsive to sensory stimulation (Dykes and Lamour 1988a, 1988b). A number of recent studies confirm that across multiple sensory modalities, neuronal activity is indeed sparse, especially in the supragranular layers of cortex (Olshausen and Field 2004, Shoham et al. 2006, Barth and Poulet 2012, Petersen and Crochet 2013). In the primary vibrissal somatosensory cortex, vS1, only a small fraction (~10%) of neurons generate action potentials in response to whisker stimulation under various experimental conditions (O’Connor et al. 2010b; Barth and Poulet 2012; Clancy et al. 2015; Peron et al. 2015). This is surprising given that the supragranular vS1 is highly connected to other cortical areas (Chen et al. 2015), and that these reciprocal connections with higher sensory areas seem crucial for whisker-mediated perceptual decisions (Yang et al. 2015; Kwon et al. 2016).

A number of factors may lead to the observation of an apparently sparse activity in vS1 (Barth and Poulet 2012; Petersen and Crochet 2013). These include anesthesia (Greenberg et al. 2008; Vincis et al. 2012; Haider et al. 2013), cortical state (Sakata and Harris 2012), as well as learning and habituation (Gdalyahu et al. 2012; Kato et al. 2015). Sparseness may also be the result of using systematically weak or simplified stimuli for precise control of stimulation parameters (Barth and Poulet 2012; Spanne and Jörntell 2015; Ranjbar-Slamloo and Arabzadeh 2017). However, a number of recent studies found that a large fraction of neurons in supragranular vS1 remain silent in both anesthetized and awake conditions or with multi-whisker, spatiotemporally complex stimuli under light anesthesia (O’Connor et al. 2010b; Crochet et al. 2011; Ramirez et al. 2014; Peron et al. 2015; Estebanez et al. 2016). *In vitro* studies show that strong translaminar (L4-to-L2/3) excitatory-excitatory connections are rare (Lefort et al. 2009) and weaker than excitatory-inhibitory connections (Helmstaedter et al. 2008). Therefore only those neurons which receive stronger excitatory input or weaker inhibition can be driven by the sensory input (Crochet et al. 2011; Petersen and Crochet 2013; Elstrott et al. 2014). These observations suggest that a small and stable fraction of neurons in L2/3 vS1 represent whisker movements while the remaining majority do not directly contribute to sensory coding (Barth and Poulet 2012; Ramirez et al. 2014). On the other hand, strong feature selectivity may underlie the sparse coding in L2/3 vS1 neurons similar to other sensory areas such as the visual and auditory cortex (Petersen and Crochet 2013). This idea is supported by the presence of high-amplitude post-synaptic potentials in most vS1 L2/3 neurons (Moore and Nelson 1998; Crochet and Petersen 2006; Poulet and Petersen 2008; Crochet et al. 2011; Ranjbar-Slamloo and Arabzadeh 2017) and their complex tuning properties (Andermann and Moore 2006; Kremer et al. 2011; Estebanez et al. 2016).

Certain gating mechanisms may allow the silent vS1 neurons to fire under special circumstances. For example, “propagation of synchrony” from L4 to L2/3 (Bruno 2011) may evoke a large fraction of neurons which would in turn effectively communicate strong, salient stimuli to downstream targets. Our previous work showed that under anesthesia, a majority of vS1 L2/3 neurons (>70%) fire action potentials in response to a high-velocity “sharp” stimulus (Ranjbar-Slamloo and Arabzadeh 2017). Here, we quantify the extent of sparse coding in the vS1 cortex of awake head-fixed mice, with loose-seal cell-attached recordings. Our data indicates that tuning to unexplored stimulus features may underlie sparse coding in vS1.

## Results

### Spiking response to piezo stimulation

We performed loose-seal cell-attached recordings in awake mice while presenting a range of whisker stimuli either applied through a piezo-electric ceramic (piezo) or by manual stimulation. We first begin by characterizing the response of vS1 neurons to piezo stimulation, which included a sharp, high-velocity stimulus, (with a peak velocity of 3.8 degree/ms) and a standard single deflection stimulus (1.2 degree/ms) presented to the neuron’s principal whisker. In the absence of anesthesia, on many trials the sharp stimulus produced a bilateral reflexive movement of the whiskers (Figure 1*C*&*D*). We therefore limited our analysis to a 30-ms post-stimulus window to quantify the response evoked by the piezo movement eliminating any potential contamination by the reflex. The choice of 30-ms window was based on the observation that the whisker reflex had a delay of >20 ms, which could in turn influence the cortical activity only after an additional ~10 ms delay (Figure 1*C*).

**Figure 1:**
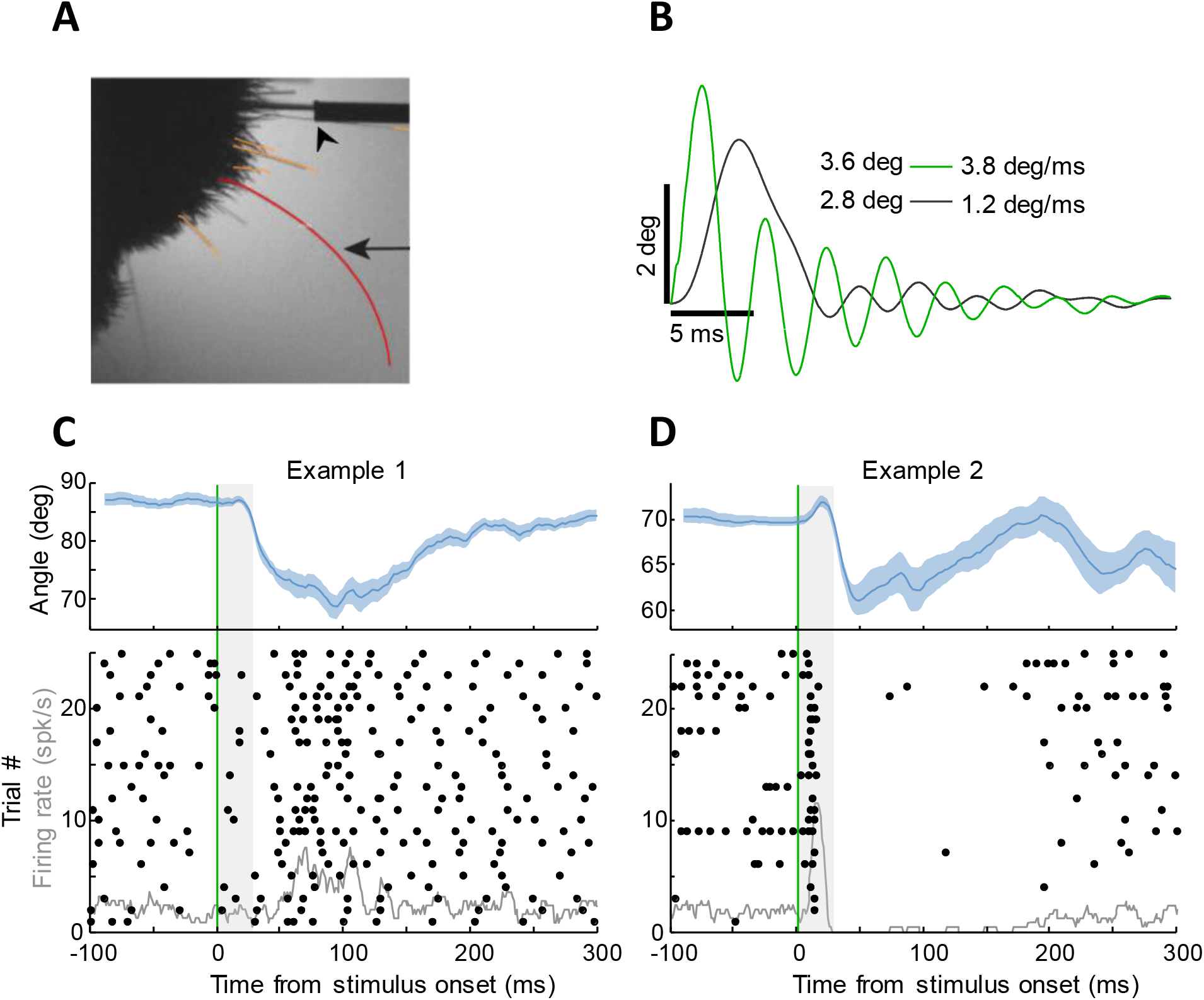
Awake recordings with piezo stimulation. ***A**: Top view of the snout showing the position of the stimulated whisker in the cannula (arrowhead). The arrow points to the tracked whisker (red). **B**: The profile of the piezo movement for the sharp (green) and standard (black) stimuli as measured by the optical sensor. Peak amplitude in degree (deg) and peak velocity in degree/ms (deg/ms) are reported for each stimulus. **C**: Top; average angle of the tracked whisker (mean ± SE) across 25 trials, aligned to the onset of the sharp stimulus (the vertical green line). Gray shading represents the 30-ms interval following the stimulus onset in which piezo-driven response is quantified. Bottom; the raster plot (dots) and the overlaid peri-stimulus time histogram (PSTH, gray trace) for the recorded neuron. **D**: As in **C** but for a second example neuron.*

For the piezo-electric stimulation protocol, a total of 155 neurons were recorded from 5 mice at various depths of the vS1 cortex (see Supplementary Figure s1*A* for details of spike waveforms, spiking raster-plots, and the classification into fast spiking (FS) and regular spiking (RS) neurons). Across all 155 neurons the average evoked response was 0.28 APs per trial for the standard stimulus and 0.44 APs per trial for the sharp stimulus (Figure 2*A*). After excluding 17 FS neurons, the average evoked response dropped to 0.13 and 0.24 APs per trial for the standard and the sharp stimuli respectively (Figure 2*B*). For FS neurons the corresponding average responses were 1.46 and 2.04 APs per trial (Figure 2*C*).

**Figure 2:**
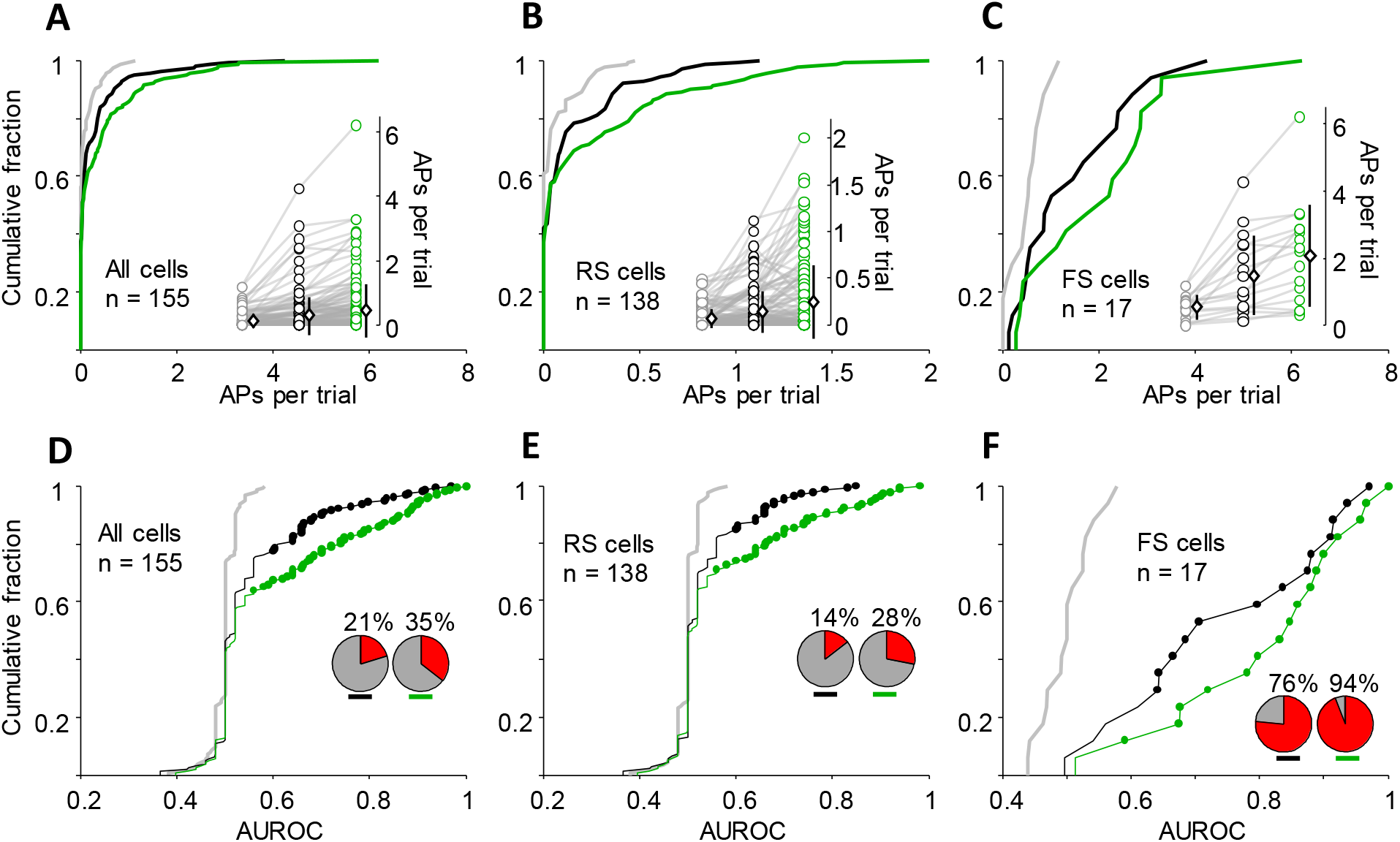
Evoked spiking activity in response to the standard and the sharp stimulus. ***A-C**: Cumulative distribution of response for all cells (**A**), and separately for RS cells (**B**) and FS cells (**C**). The cumulative distributions are plotted for the spontaneous activity (gray), and for the evoked response to the standard stimulus (black) and the sharp stimulus (green). Inset plots represent spontaneous (gray) and evoked (black and green) activities where each circle is a stimulus-neuron pair. Diamond symbols represent average (± SD) spike count across neurons. **D-F**: Cumulative distribution of the area under ROC curves (AUROC) are plotted with the same color convention as above. Filled circles indicate neurons with significant responses (p < 0.05) based on a bootstrap permutation test on the ROC analysis. Inset pie charts represent the fraction of significant AUROCs for the standard (left pie-chart) and the sharp (right pie-chart) stimulus.*

Overall, the standard stimulus resulted in significant response in 21% of neurons while the sharp stimulus evoked significant response in 35% of neurons (Figure 2*D*, ROC analysis followed by permutation test, p < 0.05). Among 138 RS neurons, 28% produced a statistically significant response to the sharp, high-velocity stimulus and this was 14% for the standard stimulus (Figure 2*E*). Most of the FS neurons were responsive to both stimuli (76% to the standard and 94% to the sharp stimulus; Figure 2*F*).

Cumulative plots in Figure 2*B* (black and green) depict heavy-tailed distributions of responses, a signature of sparseness (Olshausen and Field 2004). To quantify the degree of sparseness, we calculated population sparseness (see Equation 1 and 2 in Methods) during spontaneous activity and for the standard and the sharp stimuli presented in both anesthetized and awake conditions. Figure 3*A* illustrates population sparseness values calculated separately for the RS and FS neurons. For the RS cells recorded under anesthesia, sparseness dropped from 0.93 for stimulus zero to 0.75 for the standard stimulus and then to 0.31 for the sharp stimulus, indicating a dense activation as reported earlier (Ranjbar-Slamloo and Arabzadeh 2017). In awake condition however, sparseness remained highly stable and did not change with stimulation (0.78, 0.73 and 0.73 for the zero, the standard and the sharp stimulus respectively). As expected, FS neurons exhibited consistently low degrees of sparseness (≤ 0.40) both under anesthesia and in awake condition (Figure 3*A*). There was a negative correlation between the degree of sparseness and the fraction of responsive neurons (Figure 3*B*, Spearman’s correlation r_s_ = −0.88, p = 0.007). However, we did not always see a decrease in sparseness as the fraction of the responsive neurons increased; in awake condition, the fraction of responsive RS neurons was 14% (standard stimulus) and 28% (sharp stimulus), whereas their corresponding sparseness was almost equal (filled red circles in Figure 3*B*).

**Figure 3:**
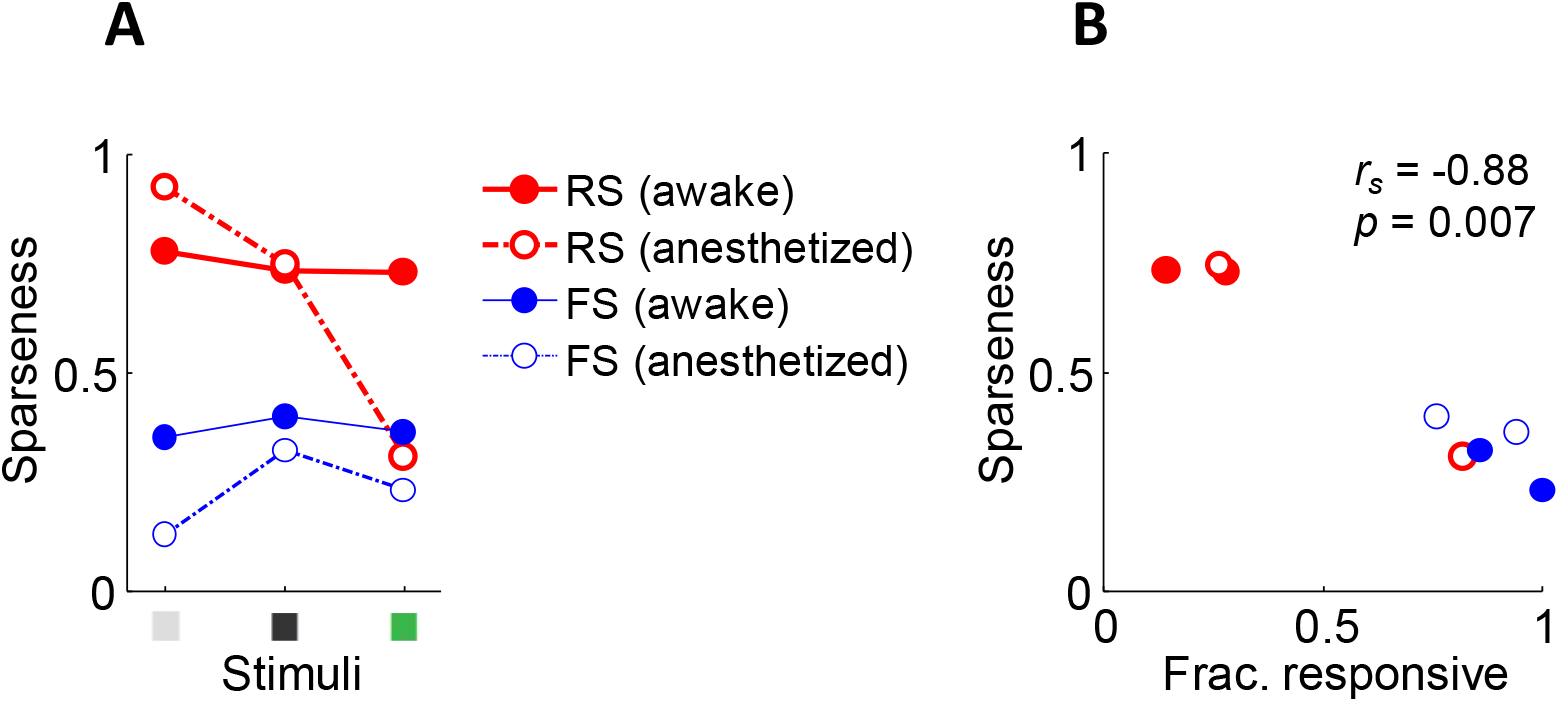
Quantification of sparseness across different states and levels of stimulation. ***A**: Spontaneous (stimulus zero), standard and sharp stimuli are represented as gray, black and green squares respectively. Filled circles with solid lines plot values for awake condition and open circles with dashed line represent values for anesthetized condition. **B**: Scatter plot of sparseness versus fraction of responsive neurons (Frac. responsive) for the standard and the sharp stimuli. Colors and symbols are preserved from panel **A**. Inset represents the Spearman’s correlation coefficient (r_s_) and its p value.*

Next, we plotted the neuronal response as a function of depth. The percentage of responsive RS neurons was similar across two depth ranges (Figure s2*A*). Across 79 RS neurons recorded in the range of 105-400 μm, 16% were responsive to the standard stimulus and 29% were responsive to the sharp stimulus (Figure s2*C*). Across 59 RS neurons recorded in the range of 403-768 μm, 12% and 27% were responsive to the standard and the sharp stimulus respectively.

### Spiking response to manual stimulation

To examine whether the observed sparseness was due to the choice of stimulus, we employed a broad range of manual stimulation. For 18 neurons, we maintained the stable recording after the termination of piezo stimulation protocol, which allowed subsequent testing with manual stimulation. For these neurons, the piezo was removed and the whisker was manually deflected. Out of 18 neurons tested for both piezo and manual stimulation, 14 were RS and 4 FS. All 14 RS neurons were responsive to at least one form of manual stimulation. Among these neurons, 64% (9 neurons) were only responsive to the manual stimulation and not to the piezo (neither standard nor sharp stimulus) while the remaining 36% (5 out of 14) were responsive to both piezo and manual stimulation. As expected, all FS neurons were responsive to both piezo and manual stimulation (n = 4).

This finding encouraged us to undertake more manual stimulation experiments and determine responsiveness in a bigger population of neurons (n = 118, 4 mice). For manual stimulation, we kept all of the whiskers intact and explored multiple ways of whisker-object contact to find the effective stimulus. This was facilitated by flexibility of the manual probing and an audio feedback, which helped to identify and repeat touch incidents that elicited spiking activity.

To identify mechanical variables underlying the neuronal response during manual stimulation, in one of the subjects we trimmed all of the whiskers except one row and tracked whiskers using high-speed videos. Figure 4 plots the mechanical variables underlying activity of three example neurons. Panel *A* illustrates an example neuron for which the spiking activity is overlaid on whisker curvature and velocity. This neuron was responsive to tapping on whisker C1 and was silent otherwise, even when the C1 whisker was pushed and released, producing high-velocity events or large curvature events (Figure 4*A* and supplementary video 1). Spike triggered average curvature (STAC) for this example neuron shows a trough preceding the spike which is followed by a sharp increase in curvature at the spike time (Figure 5*A*).

**Figure 4:**
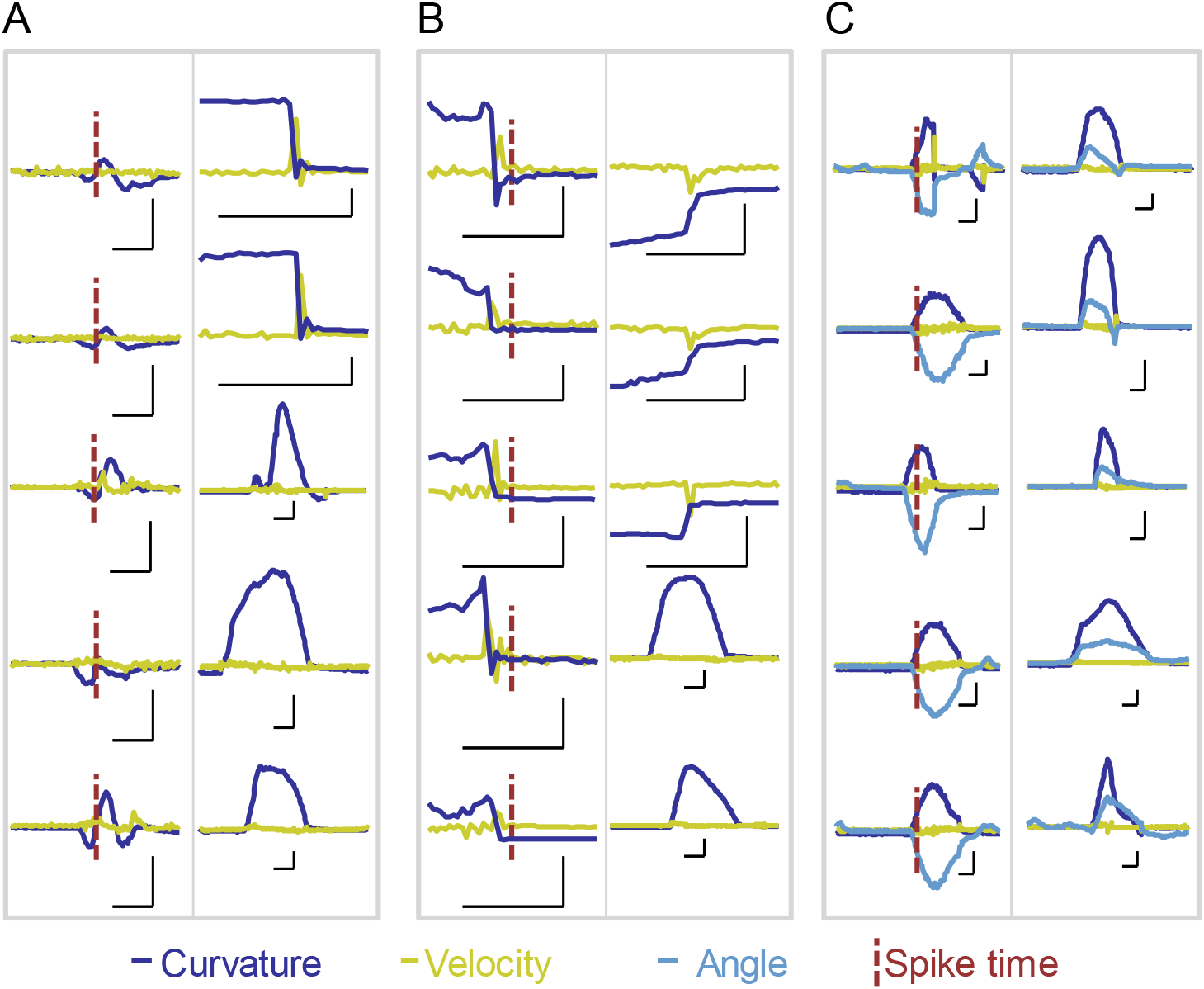
Examples of responses evoked by complex manual touch. ***A**: Example of manual whisker contacts producing reliable spiking activity (left) and those which did not produce any spiking response (right), despite large-amplitude curvature (blue) and/or velocity (lime color) events. **B**: An example neuron showing spiking response to high-velocity events (left). On the right side, the first three panels show high-velocity movements in the opposite direction, which did not elicit any spiking response. This neuron was also not responsive to high-amplitude bending of the whisker of which two examples are shown on the right. **C**: Example traces of a neuron which was responsive to rostral deflections of the principal whisker (left). Traces on the right show events produced by axial pressures which did not evoke any spikes. In all traces from **A** to **C**, vertical scale bars represent curvature (30 μm^−1^), velocity (3 degree/ms) and angle (6 degrees). Horizontal scale bars represent time (40 ms).*

**Figure 5:**
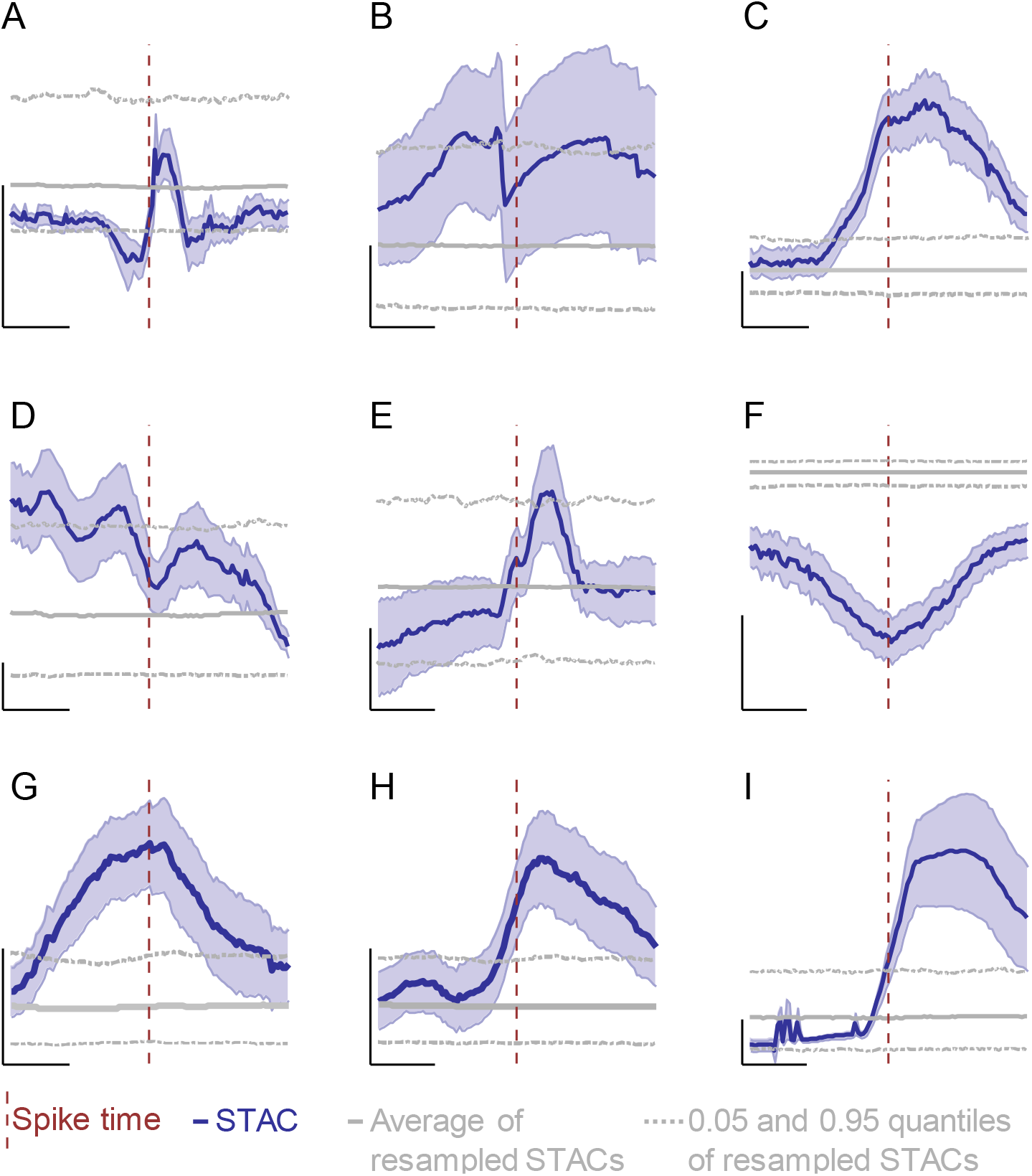
Spike triggered average curvatures (STACs) ***A-C**: STAC (dark blue, ± SE) of the example neurons shown in Figure 4 (A-C respectively). **D-I**: STACs of example neurons which were responsive to generative contact (D), tap/whisking (E), touch/whisking (F, broad tuning), lateral pressure (G), axial pressure (H) and tap/axial pressure (I). A positive curvature change indicates that the whisker tip is bent towards the rostral direction and a negative change is when the tip is bent towards the caudal direction. Solid gray lines show the average of 1000 resampled STACs. Horizontal dashed lines represent 0.05 and 0.95 quantiles of the resampled STACs. Vertical dashed lines show the spike time. Vertical scale bars represent 10 μm^−1^ curvature and horizontal scale bars represent 40 ms.*

Figure 4*B* shows another example neuron, which was exclusively responsive to high-velocity slip events produced after bending C1 whisker in rostral direction (Figure 4*B*, left). Similar slip events in the opposite direction or intense bending in axial or lateral directions did not produce any response (Figure 4*B*, right). The third example shows a neuron which was exclusively responsive to rostral bending of the whisker (Figure 4*C*, left). This neuron was not responsive to axial pressures which produced high amplitude curvature events (Figure 4*C*, right), or slip events upon release of the whisker (Figure 4*C*, left, topmost).

We recorded three neurons with exclusive response to touch during whisking. Supplementary video 2 and Figure s3*A*&*B* show an example neuron with exclusive response to contact during active touch. Whisker protractions are marked by brief troughs within the elevated curvature event where trains of spikes occurred (supplementary video 2 and Figure s3*A*&*B*). Panel *C* in Figure s3 shows a passive touch epoch marked with prominent changes in the curvature and high-velocity events. There were no responses to curvature events when the whisker was passively deflected by the object (Figure s3*C*). The occurrence of spikes in the trough of the elevated curvature is better quantified in the spike triggered average curvature (Figure 5*D*).

Figure 5 illustrates for 9 example neurons tuning to a particular stimulus category as captured by their STAC profile. This figure includes neurons with exclusive responses (Figure 4 and Figure 5 *A-C*) and neurons with broader tuning (e.g. tuning to both tapping and axial pressure (Figure 5*I*), to both tapping and whisking (Figure 5*E*) and a neuron which responded to any contact and also whisking (Figure 5*F*). The tap response was associated with a sudden change in curvature just before the spike generation (~15ms), indicating that the initial force on the whisker was effective in driving the neuron. Such tuning to a fast change in curvature was also observed for a few slip-tuned neurons (see Figure 5*B* as an example). Post-hoc analysis of STAC shows that the neuron spiked on average ~9 ms after a slip event (Figure 5*B*). Some neurons were not temporally aligned to a specific phase of the curvature events (such as those in Figure 5*F*&*G*). This was often associated with responsiveness to multiple types of contact during the experiment. Overall, these observations indicate that the curvature events can only partially capture the specific tuning of a neuron.

From the whole population of neurons recorded during manual stimulation (n = 118), we identified 11 FS and 107 RS cells (Table 1). Regardless of their tuning characteristics, all FS neurons and 78% of the RS neurons generated action potentials during sensory stimulation (i.e. a reproducible evoked response during whisking, touch or application of air puff).

**Table 1:** *Neurons tested for a response to manual stimulation*

| | | All depths | Shallow ($\leq 400$) |
| --- | --- | --- | --- |
| All types | # of cells | 118 | 65 |
|  | # responsive | 92 | 54 |
|  | % responsive | 78 | 83 |
| RS | # of cells | 107 | 61 |
|  | # responsive | 83 | 51 |
|  | % responsive | 78 | 84 |

Figure s4*B* shows that the RS neurons were almost evenly sampled from all depths and likewise responsive/nonresponsive neurons were not clustered at any specific depth range. Additionally, when we only considered neurons recorded at a depth shallower than 400 μm, 84% of the regular spiking neurons were responsive (Table 1). Further limiting the recording depth to <250 μm resulted in a similar fraction of responsive neurons (85%). Therefor most neurons, regardless of their depth, could be activated by at least one type of whisker stimulation.

Table s1 represents our classification of neuronal responses based on input category and tuning. Our input category identifies broad features, such as the receptive field size (single- or multi-whisker responses), movement source (touch or whisking), and movement direction. However, tuning identifies more specific features branching from input categories. We found 8 neurons with distinguishable response to both whisking and touch, 1 neuron with exclusive response to whisking and 81 neurons that were responsive to touch. For a large number of touch responsive neurons, we further identified a specific tuning. These include neurons responsive to lateral push but not to release or slip movement (n = 6, Table s1, movement type) and neurons exclusively responsive to slip movement but not to other types of movement (n = 5; Table s1). Notably, we found that several neurons only reliably responded to gentle touch but not to strong deflections of the whiskers (n = 11, Table s1, intensity). Such movement specific responses indicate the diverse range of stimulus parameters captured by the vS1 neurons.

## Discussion

It is widely reported that a vast majority of vS1 neurons especially in the superficial layers are silent under various experimental conditions (Dykes and Lamour 1988a; Barth and Poulet 2012; Petersen and Crochet 2013; Ramirez et al. 2014; Peron et al. 2015). An unresolved question is whether the large fraction of silent neurons contribute to coding of various stimulus features, as is the case in the visual cortex (Ohki et al. 2005), or whether they are not wired strongly enough with bottom-up excitatory input to do so. Previously, we found that under urethane anesthesia, a high-velocity sharp stimulus evoked reliable activity in a majority of L2/3 vS1 neurons (Ranjbar-Slamloo and Arabzadeh 2017). Here, we tested responsiveness of vS1 neurons in awake mice using (i) simple well-controlled piezo stimulation and cell-attached recordings (n=155 neurons, 5 mice) and (ii) manual probing of whiskers (n=118 neurons, 4 mice), which is a better representative of complex natural interactions between whiskers and objects. We found that a majority of vS1 neurons, which did not produce spiking activity in response to intense piezo-controlled stimuli, were nevertheless responsive to a specific feature or a combination of features of manual stimulation.

To determine the extent of neuronal activation by a high-velocity stimulus in awake condition, we presented a standard (1.2 degree/ms) and a sharp (3.8 degree/ms) stimulus to the principal whisker as in our previous study (Ranjbar-Slamloo and Arabzadeh 2017). Here, we reduced the number of stimuli to 2 key deflection intensities which were sufficient to observe a transition from sparse to dense activation in our anesthetized study (Ranjbar-Slamloo and Arabzadeh 2017). All other stimulus parameters were kept as in the previous anesthetized study. In contrast to the anesthetized recordings where the sharp stimulus evoked spiking activity in 74% of neurons (single whisker experiment), under awake condition, the sharp stimulus was only effective in 28% of RS neurons. To further investigate responsiveness, in a subset of neurons we tested both piezo and manual stimulation and observed that manual stimulation of the same whisker effectively drove all the tested neurons including the high fraction which did not respond to the piezo (9 out of 14). Next, we employed whisker tracking along with cell-attached recording in another subset of neurons to identify mechanical variables of the whisker-object contact during manual stimulation. We found that the neuronal tunings were only partially reflected in the spike-triggered whisker curvature (Figure 5). Overall, we found that unlike piezo stimulation, most neurons (84%) were driven by one or multiple forms of manually optimized whisker contacts. Using this approach, we also found a number of neurons to be highly selective for a certain stimulus feature and condition (e.g. generative contacts or light taps on the whisker).

For response categorization, specific tunings such as “tap only” and “slip only” were assigned to a neuron only when it was rigorously tested for various methods of stimulation including all other types of stimuli in all input categories (Table s1). Although the classifications used in the Table s1 are descriptive, tunings listed were only assigned to a neuron if the response was reliably reproduced within the identified feature. To obtain a quantitative criterion, 3-dimentional dynamics of the intact whiskers must be recorded. We were not able to measure whisker dynamics when all whiskers were kept intact (for most of the neurons). In the case of example neurons, only a row of whiskers was imaged, and thus the responsiveness could be examined only for a limited set of whiskers.

Stimulus representation in L2/3 of vS1 is known to be sparse, engaging only a small fraction of the principal neurons (Shoham et al. 2006; Barth and Poulet 2012; Petersen and Crochet 2013). Multiple scenarios which can result in the apparent sparseness of cortical area were outlined by Barth and Poulet (Barth and Poulet 2012). These include uncorrelated trial-to-trial response variability, dominance of activity in a stable population, state dependent excitability and high stimulus selectivity (Barth and Poulet 2012). Additionally, the behavioral relevance of a stimulus can influence the sparseness of the sensory representations, as does behavioral training (Gdalyahu et al. 2012; Kato et al. 2015). All these scenarios can to some extent affect the apparent sparseness in various cortical areas. For example in the visual cortex, the apparent sparseness is due to the high degree of selectivity of neurons (Ohki et al. 2005).

In rodent vS1, it is not entirely clear how the diversity of feature selectivity contributes to the widely reported sparseness. Most previous studies confined the stimulus to a single whisker with simple mechanical parameters (such as curvature, velocity or deflection amplitude). However, a few studies have explored the neuronal tuning to a complex, multi-whisker stimulation through independent piezo-electric actuators under light anesthesia (Estebanez et al. 2012, 2016; Ramirez et al. 2014). Using whole-cell recording, Ramirez et al. found that spatiotemporally complex stimuli delivered to an array of 8 whiskers did not drive spiking activity in a majority of vS1 L2/3 neurons with the average spiking responses remaining below 0.1 spikes/s for the optimized stimulus (Ramirez et al. 2014). The authors thus concluded that reports of sparseness in L2/3 is unlikely to be due to impoverished stimulation. However, in a recent study, Estebnez et al. showed that a complex multi-whisker stimulation paradigm (24 whiskers) markedly increased the fraction of responsive neurons in L2/3 such that the overall fraction of responsive neurons reached 41% (Estebanez et al. 2016). This is still much lower than the fraction of contributing neurons in an analogous study in rat visual cortex, where 75% of the L2/3 neurons significantly responded to at least one stimulus (Ohki et al. 2005). Nevertheless, the former study showed that complex feature representation prevails in populations of L2/3 neurons in vS1 (Estebanez et al. 2016). Such stimuli are difficult to deliver in a controlled manner in awake condition. Here, by manual probing, the flexible presentation of three-dimensional movements, we found that a high fraction of vS1 neurons respond to one or multiple naturalistic stimuli.

Controlled whisker stimulation and whisker imaging during behavior has revealed that the vS1 neurons are sensitive to a number of variables upon whisker movement such as velocity/acceleration, phase, direction and whisker curvature (Simons 1978; Shoykhet et al. 2000; Arabzadeh et al. 2003, 2004; Bale and Petersen 2009; Hires et al. 2015). These features are preserved from the activity of the primary sensory afferents (Arabzadeh et al. 2005; Bale and Petersen 2009; Campagner et al. 2016; Severson et al. 2017), therefore, comprise the elementary or low-level features that cortical neurons receive. The convergence of these simple features via sensory integration across multiple whiskers (Jacob et al. 2008), and possibly multiple sensorimotor pathways (Xu et al. 2012) in L2/3 of vS1 may underlie the synthesis of higher-level features which drive some neurons (Estebanez et al. 2012, 2016; Ramirez et al. 2014). Notably, such multi-whisker integrations can form emergent features such as “apparent motion” in a certain direction (Jacob et al. 2008) or contours of the objects (Kremer et al. 2011).

Rodents can map the object surfaces by a combination of body, head and whisker movements in either generative or receptive modes (Diamond and Arabzadeh 2013). In freely moving animals, interactions of whiskers with objects occur in three dimensions. Thus, spatiotemporal patterns of whisker deflections must be able to tell the animal whether for example, the head/body approaches an object or moves away from it. Although our experiment was done in a head-fixed condition, three-dimensional movement of the objects in whisker field allowed us to identify robust responses to preferred features in several neurons. For example, we found neurons which were only driven by axial pressure at the tip of a specific whisker (Table s1) and therefore might encode the approach to an object/obstacle at a certain orientation. Tap neurons may encode the location of contact point in the receptive mode sensation while push and slip neurons can provide information about object properties such as surface texture (Von Heimendahl et al. 2007; Diamond et al. 2008; Wolfe et al. 2008; Jadhav and Feldman 2010). In our experiment, objects were manually introduced to the whiskers (mostly in the receptive mode) and therefore the features related to the generative mode (such as slip events) could not be fully assessed. Nevertheless, we were able to identify 12 neurons with a generative mode response. In mice trained in an object localization task, two different sets of L2/3 neurons were found to be activated in the generative and the receptive modes (Peron et al. 2015). Based on this, we expected to observe several neurons with pure generative response. However, we observed a high degree of overlap such that 9 out of 12 neurons also responded to receptive contacts while only 3 neurons had pure generative response (Table s1). Such discrepancy might be due to the use of anesthesia for receptive mode stimulation and/or training in the earlier report (Peron et al. 2015).

In the awake condition, a number of factors can influence the mode of operation of the cortical circuit (Sabri and Arabzadeh 2018) and in turn affect responsiveness of cortical neurons; whisker position is under motor control and occasional whisking gives rise to rich whisker kinematics even in the absence of touch. Furthermore, anesthesia can damp down inhibition (Rinberg et al. 2006; Haider et al. 2013; Cazakoff et al. 2014) and alter the tuning properties of cortical neurons (Peron et al. 2015; Durand et al. 2016). In contrast to our anesthetized results, we found that in awake condition, only a quarter of the recorded neurons responded significantly to the sharp stimulus delivered by piezo. However, when we further tested the responsiveness by removing the piezo and manually deflecting the whiskers, most of these neurons were clearly tuned to often non-classical stimulus feature such as light tap, axial pressure, and generative contact. This can be attributed to a high degree of freedom in the stimulus space and/or state dependent modulation of responsiveness and tuning properties in awake condition.

Overall, our results reflect a marked difference in sparseness of vS1 regular spiking neurons between anesthetized and awake conditions. The diversity of conditions upon which vS1 neurons responded to the sensory input implies that various simple and complex features are encoded in vS1 by recruiting highly selective neurons especially in the supragranular layers.

## Methods

### Surgery

All procedures were approved by the institutional animal ethics committee. C57BL6/J were kept in individual cages and provided with ad-libitum food and water. Anesthesia was initially induced by placing the mice in a 3% isoflurane chamber before moving them to a custom-made head-restraining apparatus with a face mask continually delivering isoflurane. The skull was exposed, cleaned and covered with tissue adhesive (Vetbond; 3M, St Paul, MN) except for the area above the vS1. A custom-made aluminum head implant was then glued to the skull and secured by dental cement. The skull was then imaged using intrinsic optical imaging to map the center of the C2 barrel (Ranjbar-Slamloo and Arabzadeh 2017). A shallow well was made around the location of the C2 barrel. Cement was applied all over the skull, except the bottom of the well (~3 mm), which was covered with silicon sealant (Kwik-Kast). After the operation, the animals were given 5mg/kg of carprofen and 0.86ml/kg of penicillin (both as i.p. injection). After 3 days of recovery, the habituation to head-fixation started. During the head-fixation, the body rested in a stainless-steel tube (inner diameter of 29 mm) and the head implant was screwed to the fixation bar. Over 4 days of habituation, the duration of head-fixation was gradually increased to 1 hour. On the fifth day, the animal was anesthetized with 3% of isoflurane circulation through the face mask and the Kwik-Kast was removed. The skull was drilled circularly (Crochet 2012) over the location of C2 intrinsic signal which was marked with a shallow drill hole during the first surgery. The center of drilling (~300 μm) was thinned to the level of the dura matter, avoiding the blood vessels. The cranial window was covered with Ringer’s solution and Kwik-Kast. The animal was returned to the home cage for recovery.

### Electrophysiology

After recovery from the second surgery, the head was fixed to the apparatus while the mouse rested in the tube. The Kwik-Kast was carefully removed, and a drop of saline was applied to the craniotomy and covered the bottom of the well. The ground wire was anchored on the head-bar using plasticine and the exposed tip was placed in the well in contact with the saline. Before the pipette penetration, the dura was nicked with the tip of a 25-G needle. The pipette was filled with ACSF and advance into the brain diagonally. The angle and alignment of the pipette was set to target the center of the C2 barrel. Placement of pipette on the brain was monitored by the stereo-microscope. To identify neurons, we used two criteria; (1) the interaction of the membrane with the tip of the pipette and the subsequent increase in pipette impedance and (2) spontaneous spiking or evoked spikes by an extracellular current injection (1-10 nA).

### Whisker tracking and whisker stimulation

To facilitate whisker tracking, all whiskers were trimmed except the C row. Whiskers were illuminated form bellow by visible light. The principal whisker was inserted into a 30-G cannula which was attached to the piezo-electric ceramic and was advanced to ~4 mm from the base of the whisker. A PhotonFocus camera mounted on a Leica stereomicroscope captured high-speed videos at 400 frames per second during 1-s period around the deflection onset (0.5 s before and 0.5 s after). Frame acquisition was triggered by a National Instrument board (PCIe-6321). Stimuli were generated and delivered in dorsoventral direction as in (Ranjbar-Slamloo and Arabzadeh 2017). Two stimulus intensities were presented: the standard stimulus (2.8-degree amplitude and 1.2 degree/ms peak velocity) along with the sharp stimulus (3.6-degree amplitude and 3.8 degree/ms peak velocity, Figure 1*B*). These stimuli were presented 25 times each, in a pseudorandom order and with 1.5-2.5 s inter-stimulus interval. During manual stimulation, whiskers were imaged for 60 seconds along with the electrophysiological recording. The probes which were used for manual stimulation consisted of two standard hex keys of size 2.0 and 5.0 and a hex screw (A2-70, M5) to engage one or more whiskers. The direction and strength of the stimulus application was adjusted based on the most recent stimulation that resulted in a spiking activity of the same or previously recorded neuron. Responsiveness was defined as repeatable spike generation aligned to at least half of the repetitions. If a specific whisker-object contact did not produce a spike within a contact epoch (>5 repetitions), a different configuration was tested. This was repeated for all individual whiskers and then a combination of multiple whiskers where deflected together and different orientations where tested. Axial tuning was defined when the object moved towards the base of the whisker, deflecting the whisker by a contact with its tip. Lateral motions were applied by contacting the whisker shaft and moving it in four major directions (up, down, left and right). “Slip” response was assigned to a neuron when spiking occurred upon the release of the whisker after contact with the object. A “push” response was assigned to a neuron when the whisker contact or slip did not generate any spikes, but the spikes were fired only during large amplitude movement of the whisker. This often generated high-curvature events (Figure 4*C*). A “tap” response was defined when neurons fired in response to the object hitting the whisker with a high negative acceleration similar to a tap on a heavy object. At the end of manual probing, an air puff was applied to the whole whisker pad in rostro-caudal direction with a rubber bulb.

### Data analysis

The electrophysiological data (cell-attached) were analyzed as in our previous study (Ranjbar-Slamloo and Arabzadeh 2017). Briefly, the spikes were reliably isolated from artifacts using principal component analysis in MATLAB and spike counts per trial were calculated. We observed a fast whisker reflex (>20 ms delay) after piezo stimulus (Figure 1), which is consistent with a recently described trigeminal-facial reflexive mechanism (Bellavance et al. 2017). To isolate the piezo responses, we therefore limited the window for spike counts to 30-ms (instead of 75-ms as used previously). Average number of APs per trial was used to calculate sparseness. The measure of population sparseness (*S*) was based on the statistical methods developed by Rolls and Tovee and improved by Vinje and Gallant (Rolls and Tovee 1995; Vinje and Gallant 2000):

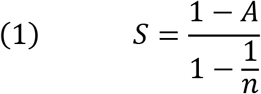

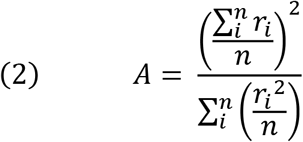

where *n* is the number of neurons and *r_i_* is the i^th^ neuron’s response (Rolls and Tovee 1995; Vinje and Gallant 2000). Receiver Operating Characteristic, ROC, analysis followed by a permutation test was used to quantify significance of the responses to the piezo stimuli at P <0.05.

Fast spiking (FS) neurons were identified based on their characteristic waveform as in our previous report (Ranjbar-Slamloo and Arabzadeh 2017). To verify the accuracy of spike waveform classification, we calculated the peak to trough of spike waveforms and plotted this measure as a function of spontaneous firing rate (Figure s1). The spontaneous firing rate of each neuron was calculated in a 500-ms interval before the stimulus onset.

For whisker tracking we used an automated software (Clack et al. 2012) to calculate whisker angle and curvature. Whisker measurements were imported to MATLAB for further analysis. Average spike triggered curvature was calculated within ±85 ms from the spike time (Figure 5). Spike times were resampled 1000 times in each recording to generate quantile boundaries for the curvature (Figure 5).

## Acknowledgements

This work was supported by the Australian Research Council (ARC) Discovery Project DP130101364 and the ARC Centre of Excellence for Integrative Brain Function CE140100007. The authors declare no competing interests.

## Supplementary figures

**Figure s1:**
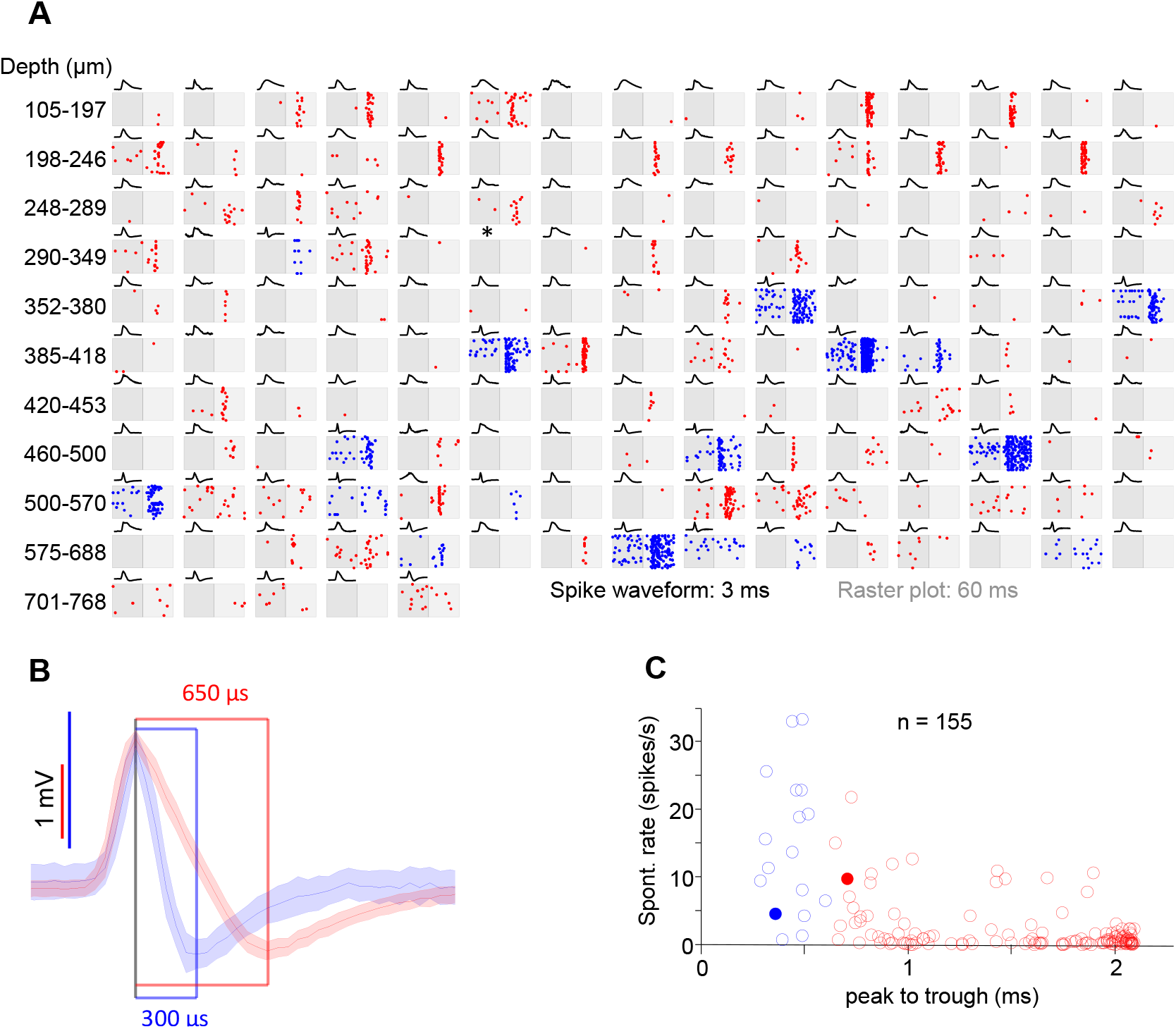
Response profile of 155 neurons recorded from 5 awake mice. ***A**: Subplots of raster plots sorted by the recording depth (from top-left to the bottom-right). Darker and lighter gray shades represent 30 ms before and 30 ms after the stimulus onset, respectively. Each dot represents the timing of a single spike relative to the stimulus onset (boundary between the two gray shades). Blue and red colored dots represent fast spiking (FS) and regular spiking (RS) neurons respectively. For each neuron the average spike waveform (normalized) is plotted on top of the spontaneous raster. For the neuron marked with an asterisk, no spike occurred during the recording protocol. **B**: Average spike waveform for two example neurons: FS (blue) and RS (red). The shaded error bands represent standard deviations. **C**: Scatter plot of spontaneous activity (y-axis) versus peak to trough (x-axis) of the average spike waveform for FS (blue) and RS (red) neurons. Filled circles represent example neurons in panel **B**.*

**Figure s2:**
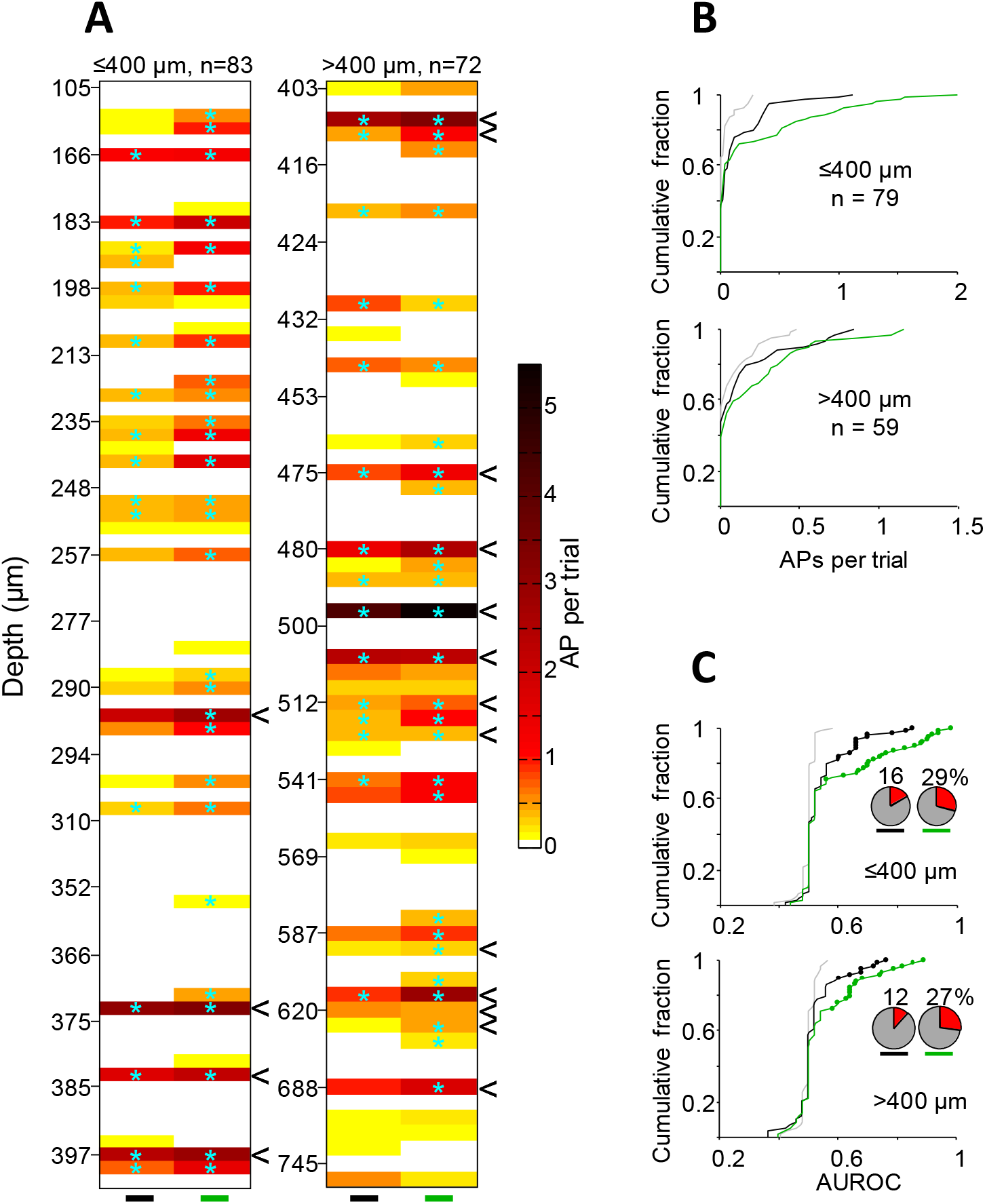
Distribution of responsive neurons across cortical depth. ***A**: The average number of evoked APs for neurons recorded at various cortical depths (n = 155). Number of APs is calculated in the 0-30 ms post stimulus onset, for both the standard (black) and the sharp (green) stimulus. “**<**“ symbols on the right edge of each plot mark fast spiking neurons. Cyan asterisks indicate significant responses based on ROC analysis (permutation test, p < 0.05). **B**: Cumulative distribution of responses in RS neurons by zero (gray), standard (black) and sharp (green) stimuli for the two depth ranges in panel **A** (≤400 μm, top; >400 μm, bottom). **C**: Cumulative distribution of the area under ROC curves (AUROCs) for the two depth ranges in panel **A**. Filled circles mark neurons with statistically significant AUROCs (permutation test, p < 0.05).Inset pie charts visualize the fraction of responsive neurons at each depth range.*

**Figure s3:**
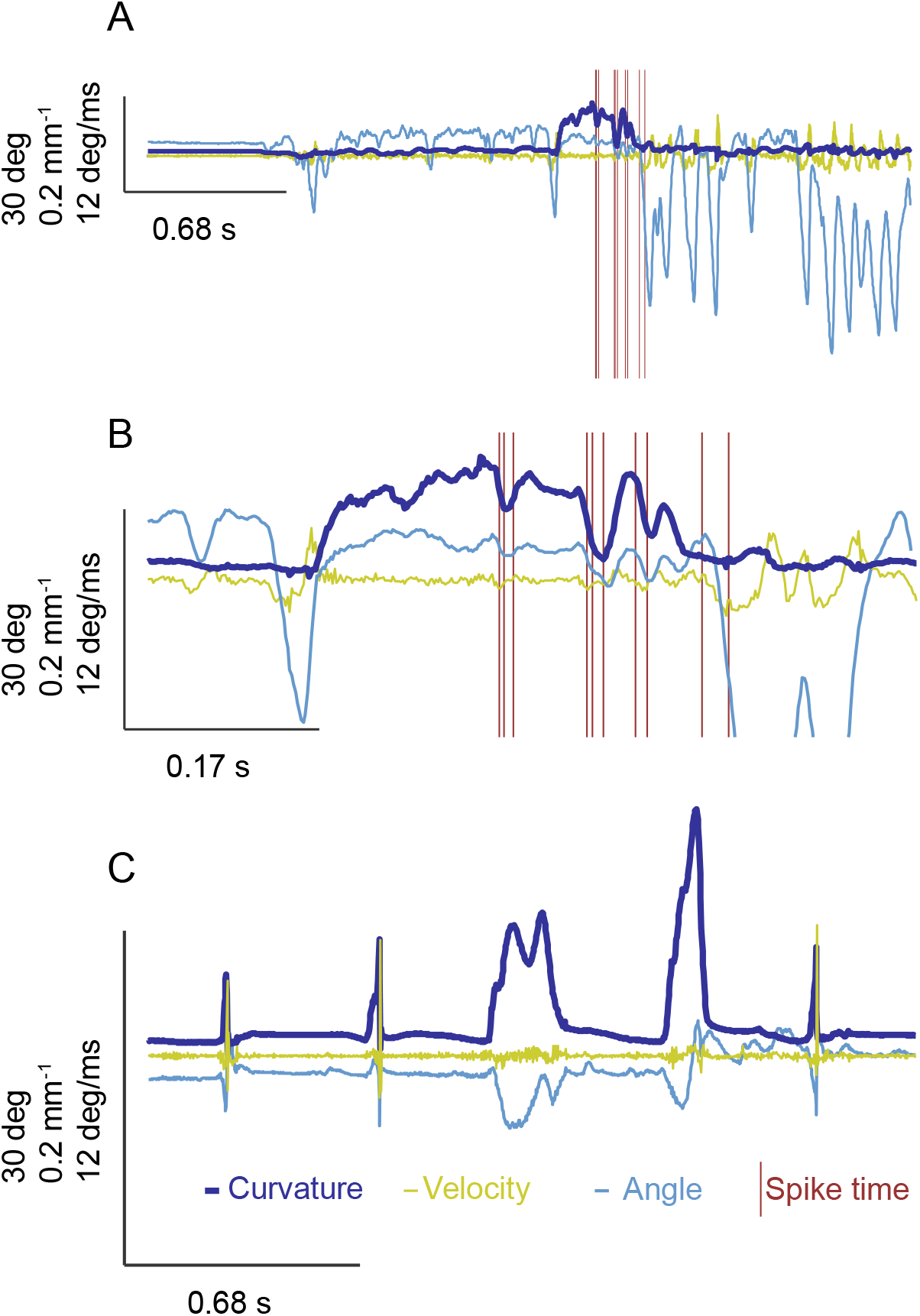
A neuron with exclusive response to generative touch (related to the Figure 5D and supplementary video 2). ***A**: Trace shows the angle (light blue) curvature (dark blue) and the velocity (lime color) of whisker C2 (also see supplementary video 2). The whisker is in a resting state and then upon whisking, the angle starts to fluctuate. The elevation in curvature marks the contact of the whisker with the object. Spikes (vertical lines in dark red) occurred during the elevated curvature event. **B**: expanded view of the elevated curvature event in panel **A**, showing that the spikes occurred during brief troughs during elevated curvature, marking the voluntary push of the whisker against the object (supplementary video 2). **C**: Passive contacts of the same whisker produced large amplitude curvature and velocity events, but did not generate any spikes.*

**Figure s4:**
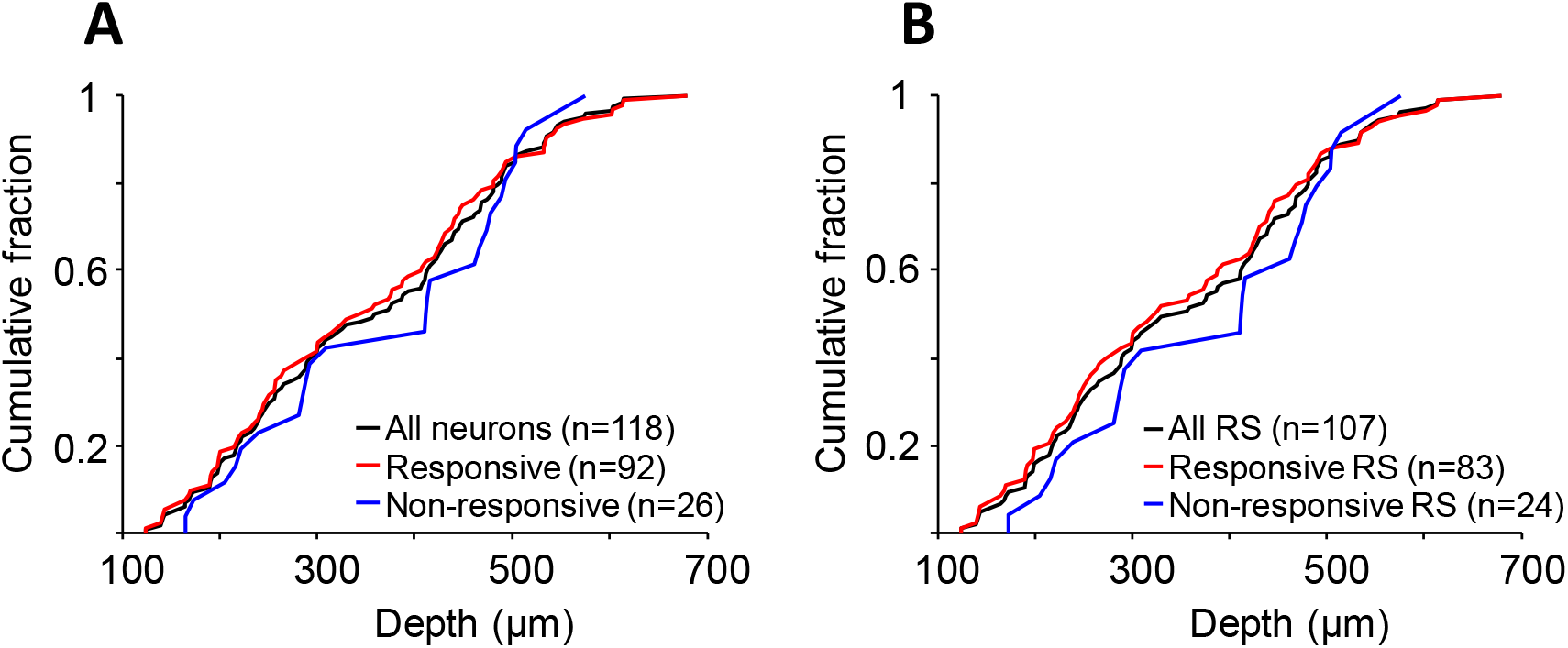
Depth distribution of the recorded neurons. ***A**: Cumulative distribution of all recorded neurons as a function of depth (black) and separately for the responsive (red) and non-responsive (blue) neurons. **B**: Same as **A** but only for the RS neurons.*

**Table s1:**
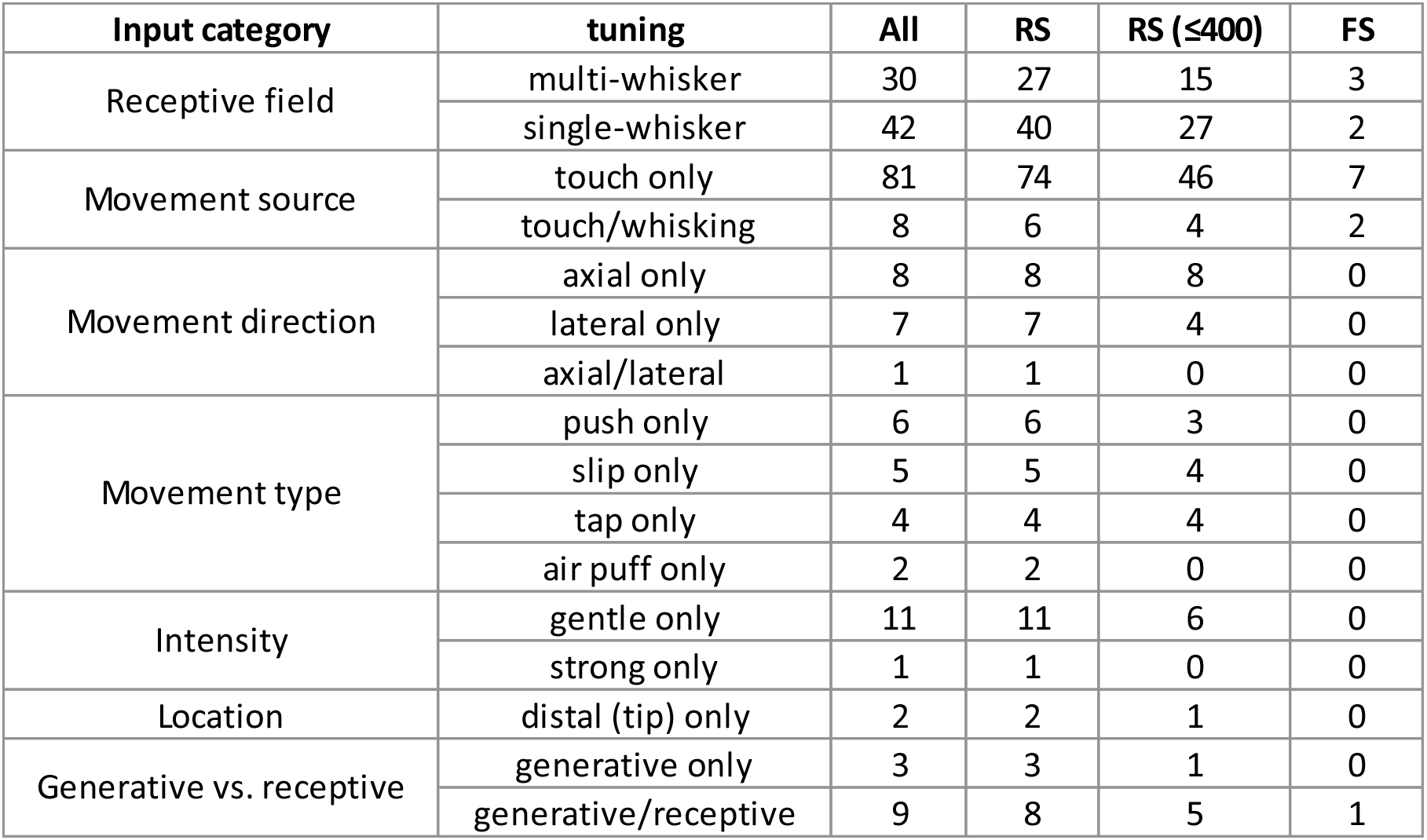
*Various tunings of responsive neurons to each class of stimuli*.

| Input category | tuning | All | RS | RS ($\leq 400$) | FS |
| --- | --- | --- | --- | --- | --- |
| Receptive field | multi-whisker | 30 | 27 | 15 | 3 |
|  | single-whisker | 42 | 40 | 27 | 2 |
| Movement source | touch only | 81 | 74 | 46 | 7 |
|  | touch/whisking | 8 | 6 | 4 | 2 |
| Movement direction | axial only | 8 | 8 | 8 | 0 |
|  | lateral only | 7 | 7 | 4 | 0 |
|  | axial/lateral | 1 | 1 | 0 | 0 |
| Movement type | push only | 6 | 6 | 3 | 0 |
|  | slip only | 5 | 5 | 4 | 0 |
|  | tap only | 4 | 4 | 4 | 0 |
|  | air puff only | 2 | 2 | 0 | 0 |
| Intensity | gentle only | 11 | 11 | 6 | 0 |
|  | strong only | 1 | 1 | 0 | 0 |
| Location | distal (tip) only | 2 | 2 | 1 | 0 |
| Generative vs. receptive | generative only | 3 | 3 | 1 | 0 |
|  | generative/receptive | 9 | 8 | 5 | 1 |

***Snapshots of supplementary videos**: snapshots of supplementary videos showing the object and the tracked whiskers just after the spiking activity of two example neurons (red dots). Traces of whisker curvature (dark blue), velocity (lime color) and angle (light blue) are overlaid on the raw image with the values for to the current frame on the far right of the traces.*

***Video 1** corresponds to the neuron in Figure 4A and 5A. **Video 2** corresponds to the neuron in Figure 4D and s3. Note that on the right panel the whisker C2 is bent due to the contact with the object way before the spiking activity.*

